# Characterization of novel regulatory modules controlling leaf angle in maize

**DOI:** 10.1101/2021.02.02.429433

**Authors:** Xianglan Wang, Xiaokun Wang, Shilei Sun, Xiaoyu Tu, Kande Lin, Lei Qin, Xingyun Wang, Gang Li, Silin Zhong, Pinghua Li

## Abstract

Leaf angle is an important agronomic trait determining maize planting density and light penetration into the canopy, and contributes significantly to the yield gain in modern maize hybrids. However, little is known about its molecular mechanism beyond the *Liguless1* (LG1) and *Liguless2* (LG2) genes. In this study, we found that transcription factor *ZmBEH1* is targeted by ZmLG2 and regulates the leaf angle formation by influencing the sclerenchyma cells layers on the adaxial side. ZmBEH1 can interact with transcription factor ZmBZR1, whose expression is directly activated by ZmLG1. Both ZmBEH1 and ZmBZR1 bind to the promoter of *ZmSCL28*, the third transcription factors that influences the leaf angle. Our study demonstrates novel regulatory modules controlling leaf angle, and provides new gene editing targets for creating optimal maize architecture suitable for dense-planting.

## Introduction

Maize yields have increased dramatically since the late 1930’s due in large part to the increased planting densities (Duvick, 2005). The selection of upright leaf architecture enables plants to be grown at higher density while minimizing the shading of neighboring plants and increasing the efficiency of light capture. Leaf angle, the trait that defined as the inclination between the leaf blade midrib and the vertical culm, contribute significantly to the upright leaf architecture. Recently, the identification of two quantitative trait loci, *UPA1* and *UPA2,* which conferred the upright plant architecture by reducing the leaf angle, and then dramatically improved planting density and enhanced grain yield (Tian et al., 2019), further support the critical role of leaf angle alteration in maize breeding.

The establish of the leaf angle in maize is determined by the ligular region, which encompasses the ligule and auricle, serves as a hinge at the blade-sheath boundary, allowing the leaf blade to project away from the stem. Two classical genes, Liguless1 (LG1), a Squamosa Promoter Binding Protein domain containing transcription factor (TF) (Moreno et al., 1997) and Liguless2 (LG2), a basic leucine zipper transcription factor family member (Harper and Freeling, 1996; Walsh et al., 1998), play key roles in the ligular region formation that altered the leaf angle. In *lg1* mutant, ligule and auricle are not formed and the plants exhibit excessively erect leaves (Moreno et al., 1997). In *lg2*, ligule and auricle are often absent or positioned incorrectly, also cause the upright plant architecture with extreme small leaf angle (Lambert and Johnson, 1978;Walsh et al., 1998; Mantilla-Perez and Salas Fernandez, 2017). Introgression of *lg2* mutant alleles into maize hybrid lines presented an increased grain yield (Pendleton et al., 1968; Lambert and Johnson, 1978), however, the extremely erect leaf limits the commercial use of this allele. Even though, ZmLG1 and ZmLG2 could provide genetic clue to optimize leaf angle for dense planting, such as identify their downstream targets. The genetic basis by which genes affect leaf angle in maize need further exploration.

Several genes in maize have been identified to control leaf angle (Muehlbauer et al., 1999; Ku et al., 2011; Zhang et al., 2014; Tian et al., 2019; Cao et al., 2020; Ren et al., 2020), and most of them are related to the phytohormone brassinosteroids (BRs), e.g., *BRD1* (brassinosteroid C-6 oxidase1) and *BRI1* (Brassinosteroid Insensitive 1). The overexpression of *BRD1*, which encodes brassinosteroid C-6 oxidase1 to catalyze the last step of brassinosteroid synthesis, increased the leaf angle by enlarging auricle and decreasing the number of sclerenchyma cells on the adaxial side; and knockout the function of *BRI1*, which encodes a Leu-rich repeat receptor kinase that responsible for BR binding to start BR signal transduction, exhibit the upright leaf with decreased auricle formation (Makarevitch et al., 2012; Tian et al., 2019).

The BR singling is send from BRI1 receptor kinase at the cell surface to the BZR1/BES1 (Brassinazole Resistant 1/ BRI1-EMS-Suppressor1) transcription factors, like BZR1 in *Arabidopsis*, which directly binds to the BR response element (BRRE, CGTGT/CG) to regulate the expression of the downstream BR-responsive genes (Li and Jin, 2007). The BZR1 family members act not only as major transcription factors in BR singling pathway, but also as mediators to participate in plant development and abiotic stress (Wang et al., 2014). In *Arabidopsis*, BZR1 promotes phloem and xylem Differentiation (Saito et al., 2018), and interacts with GATA2, a transcription factor in the light signaling pathway, to regulate hypocotyl elongation of seedlings (Luo et al., 2010). A recent study showed that anther locule development was also regulated by BZR1 (Chen et al., 2019). In addition, BZR1 serve as regulator in abiotic stress. For instance, BZR1 positively modulates plant freezing tolerance through CBF-dependent and CBF-independent pathways (Li et al., 2017). In rice, OsBZR1 was identified by its homology to *Arabidopsis* BZR1, and the Loss-of-function of *OsBZR1* reduced BR sensitivity, presented dwarfism phenotype with erect leaves, indicating the critical roles of *OsBZR1* in BR signal pathway (Bai et al., 2007). To date, very few researches focus on maize BZR transcription family and our understanding of the role of BZRs in maize is still limited.

In this study, we screened the downstream target of ZmLG2 and identified a *BZR1/BES1* homolog gene named *ZmBEH1*. The loss-of-function of *ZmBEH1* exhibits upright plant architecture. Interestingly, the maize ZmBZR1, which also influences the leaf angle, acts downstream of ZmLG1, and interacts with ZmBEH1 to co-regulate the expression of *ZmSCL28*, a rice *DLT* (Tong et al., 2009) homolog that altering the leaf angle in maize. These findings reveal a ZmLLG2/LG1-ZmBEH1/BZR1-ZmSCL28 cascades primarily regulates the leaf angle in maize, which will benefit to deepen our understanding on BR response in maize and help to optimize maize plant architecture for dense planting.

## Results

### ZmLG2 binds to the promoter of *ZmBEH1*

*ZmLG2*, which encodes a bZIP transcription factor, is a classical regulator controlling maize ligule development (Harper and Freeling, 1996; Walsh et al., 1998; Bolduc et al., 2012).

To study the regulatory mechanism mediated by ZmLG2, we searched the maize 104 TF ChIP-Seq data collection (Tu et al., 2020), and found a putative target gene named *ZmBEH1* (BZR1/BES1 homolog), which is a homolog to the rice BZR3 (Supplemental Fig. S1), has a strong binding peak of *ZmLG2* in front of TSS (Fig. 1A).

**Figure 1.**
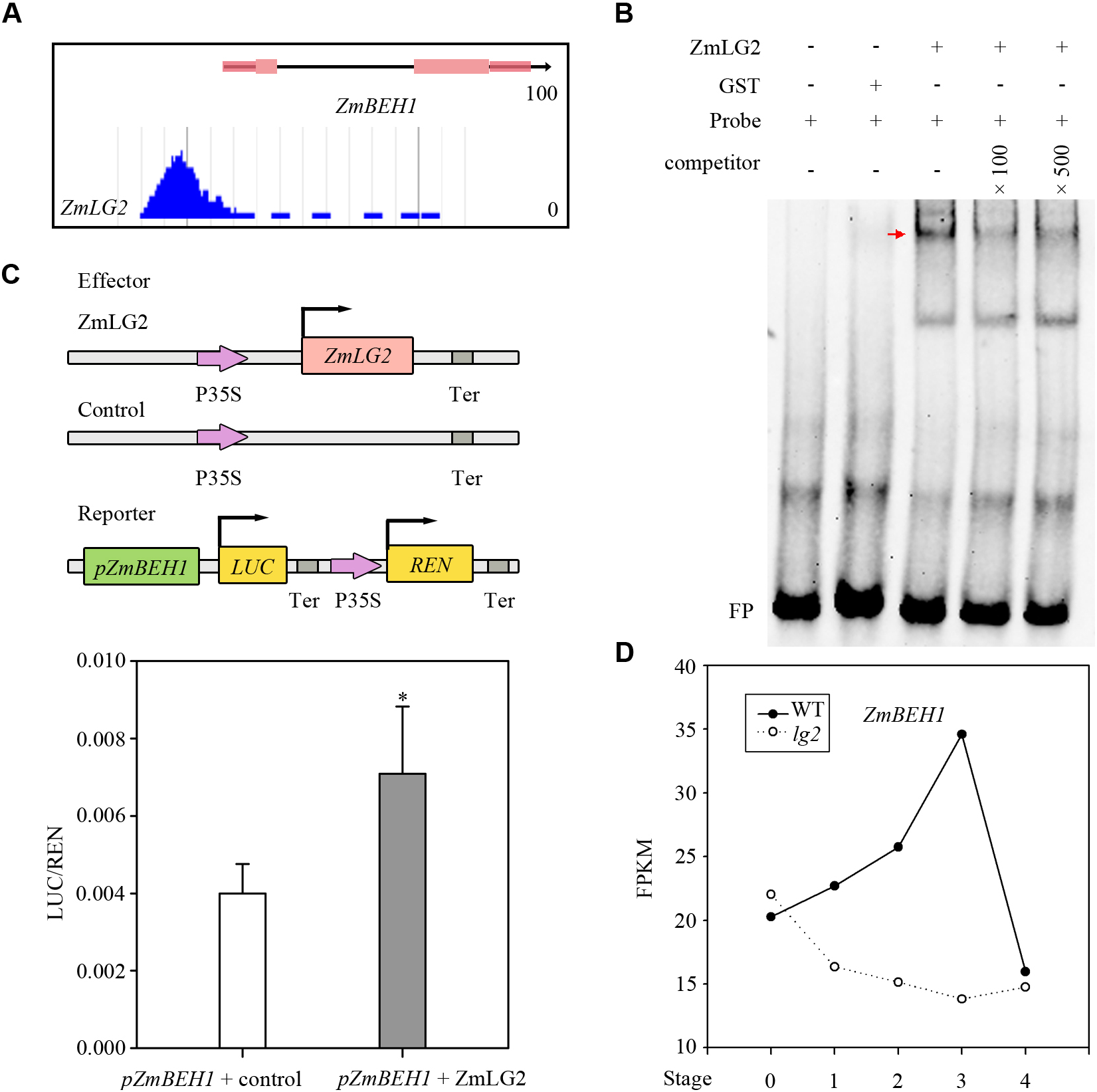
ZmLG2 directly regulates the expression of *ZmBEH1.* A, ChIP-Seq shows ZmLG2 binds to the *ZmBEH1* promoter. B, ZmLG2 directly binds to the *ZmBEH1* promoter in EMSA. “−” indicates the absence of proteins. “+” indicate the present; FP indicates free probe. C, Dual-luciferase assay shows relative transactivation of ZmLG2 to *ZmBEH1* promoter. The coding sequence of ZmLG2 driven by the 35S promoter was used as an effector, and the empty vector was used as an effector control in transient Luciferase assay performed in maize protoplasts. The vector with the *Renilla reniformis* (REN) gene driven by a 35S promoter and firefly luciferase (LUC) gene driven by promoter sequence from *ZmBEH1* were used as the reporter. The LUC/REN ratio represents the relative activity of the promoters. Error bars represent SD (n = 5) *represents P < 0.05 determined by Student’s t test. D, RNA-Seq data shows the expression profiles of *ZmBEH1* in different maturation stage of the ligular region (S0 to S4) in WT and *lg2* mutant. Details showed in Supplemental Fig. S2, A and C.

To verify the affinity between ZmLG2 and *ZmBEH1* promoter, electrophoretic mobility shift assay (EMSA) was conducted using purified recombinant ZmLG2-GST fusion protein and a synthesized 36 bp probe deriving from the promoter of *ZmBEH1* (Material and Methods). As shown in Figure 1B, ZmLG2-GST fusion protein has high affinity with the *ZmBEH1* promoter probe, however, this binding was significantly inhibited by unlabeled DNA in the competitive assay, indicating the binding of ZmLG2 to *ZmBEH1* 36 bp promoter probe is highly specific (Fig. 1B). In addition, Dual-luciferase (LUC) transient transcriptional activity assay performed in the maize protoplasts with ZmLG2 driven by the 35S promoter as an effecter and luciferase driven by the *ZmBEH1* promoter as the reporter. The result also confirmed that *ZmBEH1* promoter could be activated by ZmLG2 in vivo (Fig. 1C).

Next, we checked the expression profiles of *ZmBEH1* in *lg2* mutant with RNA-seq using ligule tissues from five different maturation stages (from ligule band initiation in S0 to auricle matured in S4, Material and methods, Supplemental Fig. S2, A and C, unpublished). The expression of *ZmBEH1* was significantly reduced in *lg2* relative to the wild type (WT) (Fig. 1D). All these results indicate that the *ZmLG2* positively regulate the expression *ZmBEH1,* and *ZmBEH1* is the direct downstream target of ZmLG2.

### ZmBEH1 Mu-insertion mutant and CRISPR lines showed reduced leaf angle

To study the function of ZmBEH1, we obtained the uniform Mu insertion mutant of *ZmBEH1* (named *beh1-1*) from maize stock center. Genomic DNA and real-time PCR results showed that the Mu transposon was inserted to the 5’-UTR of *ZmBEH1* and dramatically decreased its expression in the mutant (Fig. 2, A and B, Supplemental Fig. S3). The decreased expression of *ZmBEH1* is consistent with the slow growing phenotype of the *beh1-1* plants, with the total heights much shorter than that of the WT in both 15-day-old seedlings stage (V2 stage) and 54-day-old stage (V10 stage) (Fig. 2C). Interestingly, when the WT height stopped to increase after silking, *beh1-1* mutant could catch up. Another phenotype of *beh1-1* was its erect leaves. The leaf angle of the mutants decreased 34.9% in V2 stage and 30.9% in V10 stage compared with the WT (Fig. 2D). We noticed a smaller ligular area and margin width in the *beh1-1* than those in the WT, which decreased the auricle size at V2 and V10 stage and then decreased the leaf angle (Fig. 2E). Cross-sections of the ligular region revealed a reduced number of sclerenchyma cell layers at the adaxial blade-sheath junction in the mutant at the midrib (Fig. 2F), suggesting that in maize, leaf angle is mediated at least in part through the regulation of adaxial sclerenchyma development.

**Figure 2.**
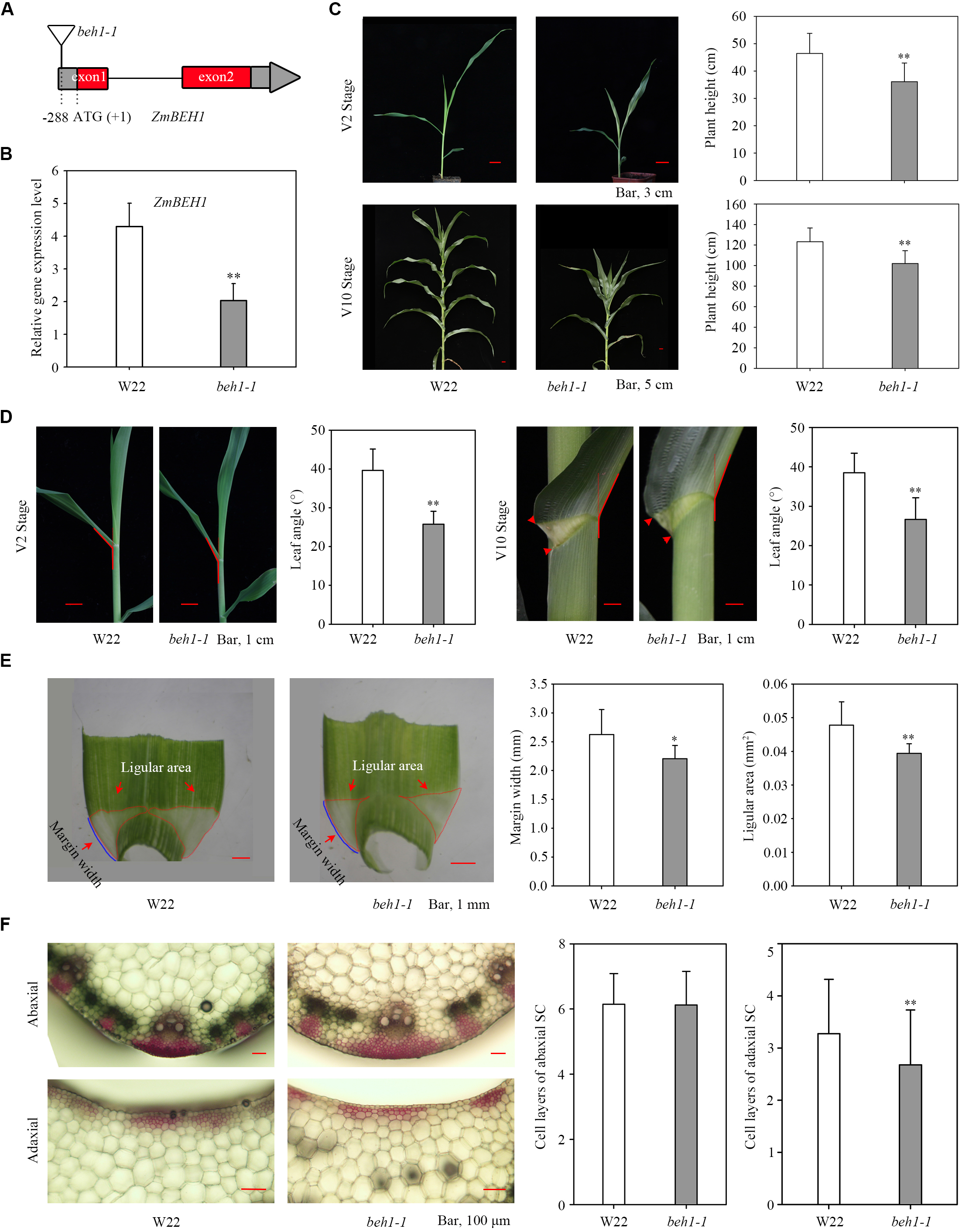
Phenotypes of *beh1-1* plants. A, Uniform Mu-mediated insertion of *ZmBEH1*. A maize transposon-insertion line *beh1-1* was obtained from the maize Uniform Mu resource, which carried a Mu insertion (UFMu-13557) in the 5’-UTR of *ZmBEH1*. B, Expression of *ZmBEH1* gene in WT inbred line W22 and *beh1-1* plants at V2 stage. **represents P < 0.01 determined by Student’s t test. C, Plant height of WT inbred line W22 and *beh1-1* plants at V2 and V10 stages. From C to E, * represents P < 0.05, **represents P < 0.01 determined by Student’s t test, n = 20. D, Leaf angle changes between WT inbred line W22 and *beh1-1* plants at V2 and V10 stages. E, Quantitative measurements of leaf ligule margin width and ligular area in WT inbred W22 and *beh1-1* plants at 15-day-old seedling stage. F, Cross-sections of the ligule from WT inbred line W22 and *beh1-1* plants at V2 stage. The sclerenchyma cell (SC) layers stained red with safranin. Number of SC cell files at the adaxial and abaxial side were calculated from 10 replicates. Error bars are SD. **P < 0.01. Scale bars, 100 μm.

To confirm the function of *ZmBEH1*, we used CRISPR/Cas9 system to edit *ZmBEH1* specifically. The two obtained CRISPR mutants were named *beh1-2* and *beh1-3*, which have a 76 bp deletion and one base pair change, respectively (Supplemental Fig. S4A). Consistent with the phenotype of *beh1-1*, *beh1-2* and *beh1-3* exhibit erect leaves, fewer layers of sclerenchyma cells on the adaxial side and shortened plant height at V2 stage (Supplemental Fig. S4, B-D). Collectively, the evidence from these lines suggest that *ZmBEH1* plays a significant role in leaf angle formation in maize.

### Yeast two-hybrid and split luciferase assays showed that ZmBEH1 interacts with ZmBZR1

In order to identify the partners that interact with ZmBEH1, we screened the B73 leaf cDNA yeast two hybrid library using ZmBEH1 as a bait. A protein that has high homology with OsBZR1 and AtBZR1/BES1 (Supplemental Fig. S1) was identified to have strong interaction with ZmBEH1 (Fig. 3A), which we named as ZmBZR1.

**Figure 3.**
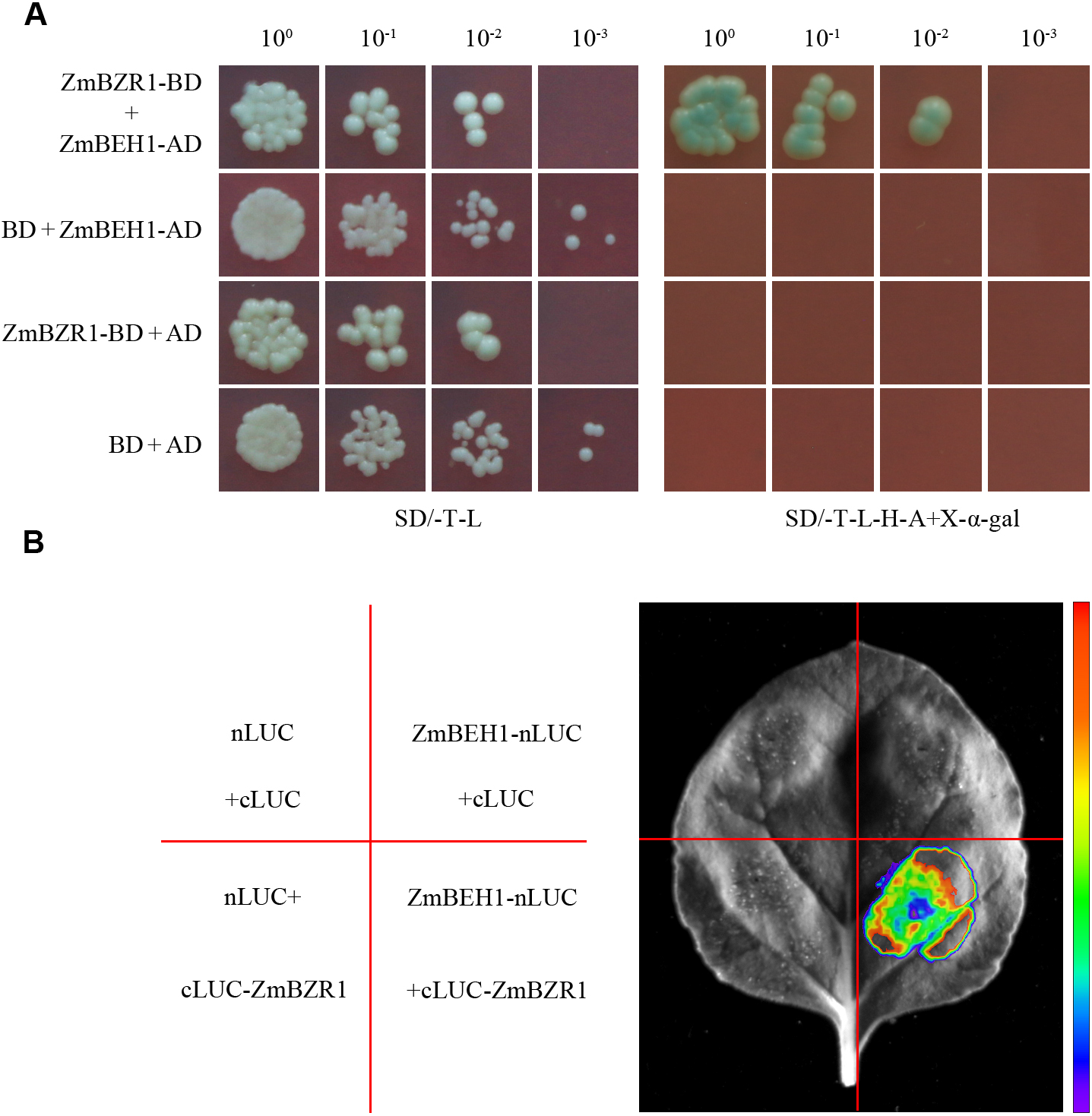
ZmBEH1 interacts with ZmBZR1. A, Interaction between ZmBZR1 and ZmBEH1 in yeast two-hybrid assays. Different concentrations of co-transformed yeast cells were spotted on synthetic dropout (SD) medium without tryptophan and leucine (SD/-T-L), or without tryptophan, leucine, histidine and adenine, and plus 20 mg/mL X-α-gal (SD/-T-L-H-A + X-α-gal). B, Interaction between ZmBZR1 and ZmBEH1 in the split luciferase assay.

To further confirm the interaction between ZmBZR1 and ZmBEH1 *in vivo*, we then performed the split luciferase assay with ZmBEH1 fused to the N terminus of LUC (ZmBEH1-nLUC) and ZmBZR1 fused to the C terminus of LUC (cLUC-ZmBZR1). When the *Agrobacterium* strain GV3101 containing both the ZmBEH1-nLUC and cLUC-ZmBZR1 constructs was injected into the tobacco leaf, a strong luciferase signal was detected, suggesting that the interaction between ZmBEH1 and ZmBZR1 also occurred *in vivo* (Fig. 3B).

### ZmBZR1 Mu-insertion mutants and CRISPR lines showed reduced leaf angle

To study its function, we obtained two potential loss-of-function mutants of ZmBZR1. One has a uniform Mu inserted to the first exon of *ZmBZR1* and the other one has Mu inserted to the first intron, which were referred to as *bzr1-1* and *bzr1-2,* respectively (Fig. 4A, Supplemental Fig. S3). We found that the expression of *ZmBZR1 in* both mutant alleles was significantly decreased (Fig. 4B), and the plants exhibit the erect leaf phenotype as *beh1-1*. Compared with the WT, the leaf angle of *bzr1-1* and *bzr1-2* decreased about 27.5% and 31.3% at V2 and V10 stage, respectively (Fig. 4D). Corresponding to this, the margin width and area of ligular region decreased in both *bzr1-1* and *bzr1-2* mutants (Fig. 4E), so does the sclerenchyma layers at adaxial side (Fig. 4F).

**Figure 4.**
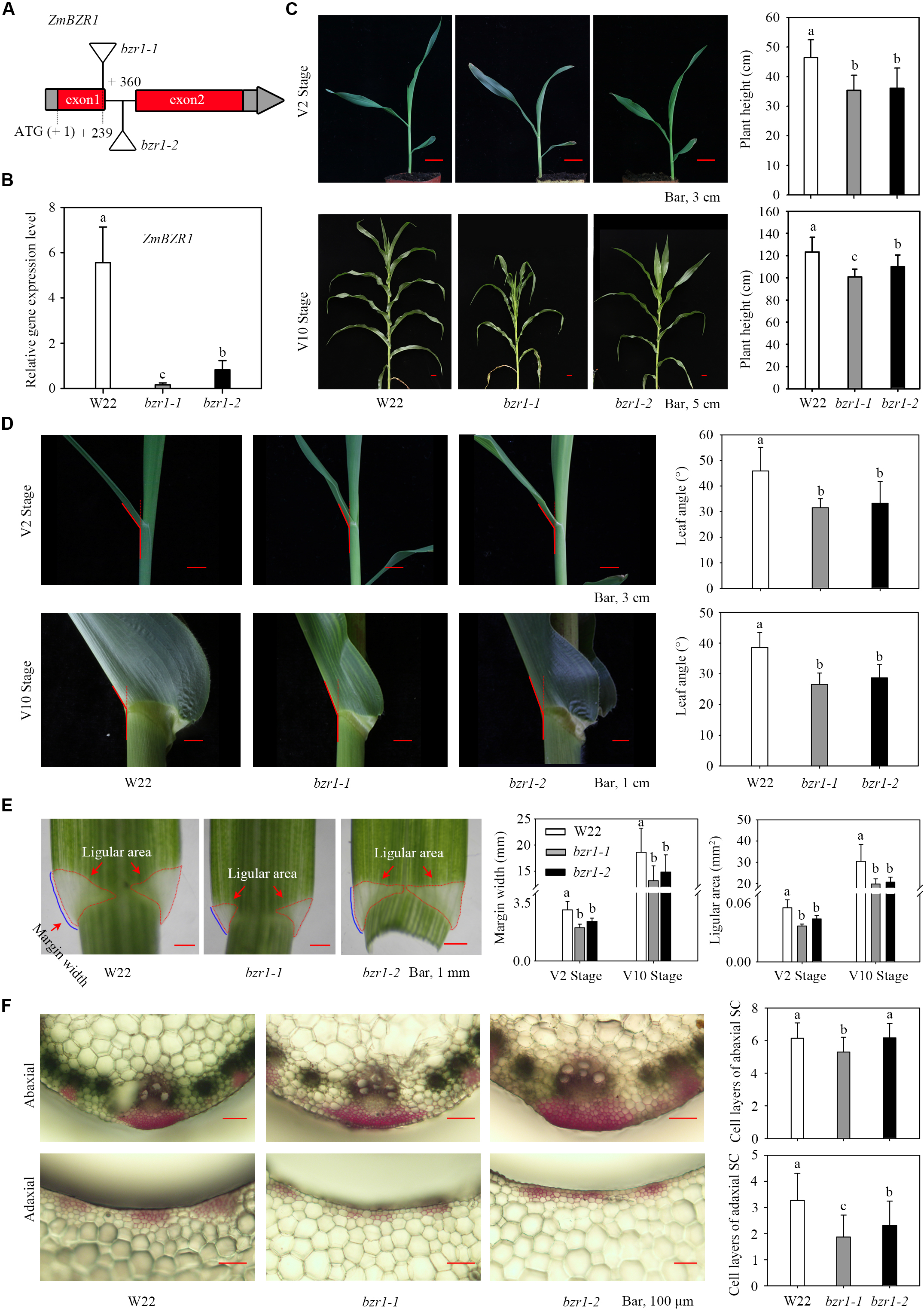
Characterization of *bzr1-1* and *bzr1-2* mutants. A, Gene model shows the inserted position of Uniform Mu (UFMu-13537 and UFMu-03258) in the 1^st^ exon (*bzr1-1*) and 1^st^ intron (*bzr1-2*) of *ZmBZR1* gene. B, Expression analysis of *ZmBZR1* in WT inbred line W22 and *bzr1-1*, *bzr1-2* plants at V2 stage. C, Plant height of WT inbred line W22 and *bzr1-1*, *bzr1-2* plants in V2 and V10 stages. Different letters above the columns indicate statistically significant differences between groups (n=20). D, Leaf angle changes between WT inbred line W22 and *bzr1-1*, *bzr1-2* plants in V2 and V10 stages (n=20). E, Quantitative measurements of leaf ligule margin width and ligular area in WT inbred line W22 and *bzr1-1*, *bzr1-2* plants at V2 and V10 stages (n=20). Different letters above the columns indicate statistically significant differences between groups. F, Cross-sections of the ligular region from WT inbred line W22 and *bzr1-1, bzr1-2* plants at V2 stage. The sclerenchyma cell (SC) layers stained red with safranin. Number of SC cell files at the adaxial and abaxial side were calculated from 10 replicates. Error bars are SD. Scale bars, 100 μm.

At V10 stage, the *bzr1-1* reached about 81.8% and *bzr1-2* reached about 89.3% of the WT height (Fig. 4C). However, when compared another nine important agronomic traits, including ear height, leaf length, leaf width and tassel branch number between *bzr1* and the WT plants at blister stage (Supplemental Fig. S5), the difference observed between *bzr1-1* and WT were much more dramatic than that in *bzr1-2*, indicating that the Mu inserted in the first intron of *ZmBZR1* caused a week allele.

In addition, another two mutant alleles of *ZmBZR1* (*bzr1-3* and *bzr1-4*) which presented by the 42 bp and 72 bp deletion in the first exon, were generated through CRISPR/Cas9 system (Supplemental Fig. S6A). The reduced leaf angle and shortened plant height were consistent with the phenotype of *bzr1-1* (Supplemental Fig. S6B).

### ZmBEH1 and ZmBZR1 act synergistically to regulate the leaf angle

We suspected whether ZmBZR1 and ZmBEH1 work redundantly in regulating the maize leaf angle. To test this, we created double mutants of *bzr1-1 beh1-1* and *bzr1-2 beh1-1,* and measured the leaf angle at V10 stage. Compared with the single mutant plants, the leaf angle in the double mutants decreased around 6°and 5° in *bzr1-1 beh1-1* and *bzr1-2 beh1*, respectively (Fig. 5A). Consistently, double mutant plants have upright leaf angle, a narrower margin width and smaller ligular area, resulting in reduced auricle size at V10 stage (Fig. 5B). Except these, the double mutants were almost identical to the single mutants in terms of other agronomic traits e.g., leaf length and width (Supplemental Fig. S7, A and B). Together with the observation that ZmBZR1 and ZmBEH1 physically interact, we propose that they might co-operate with each other to regulate leaf angle.

**Figure 5.**
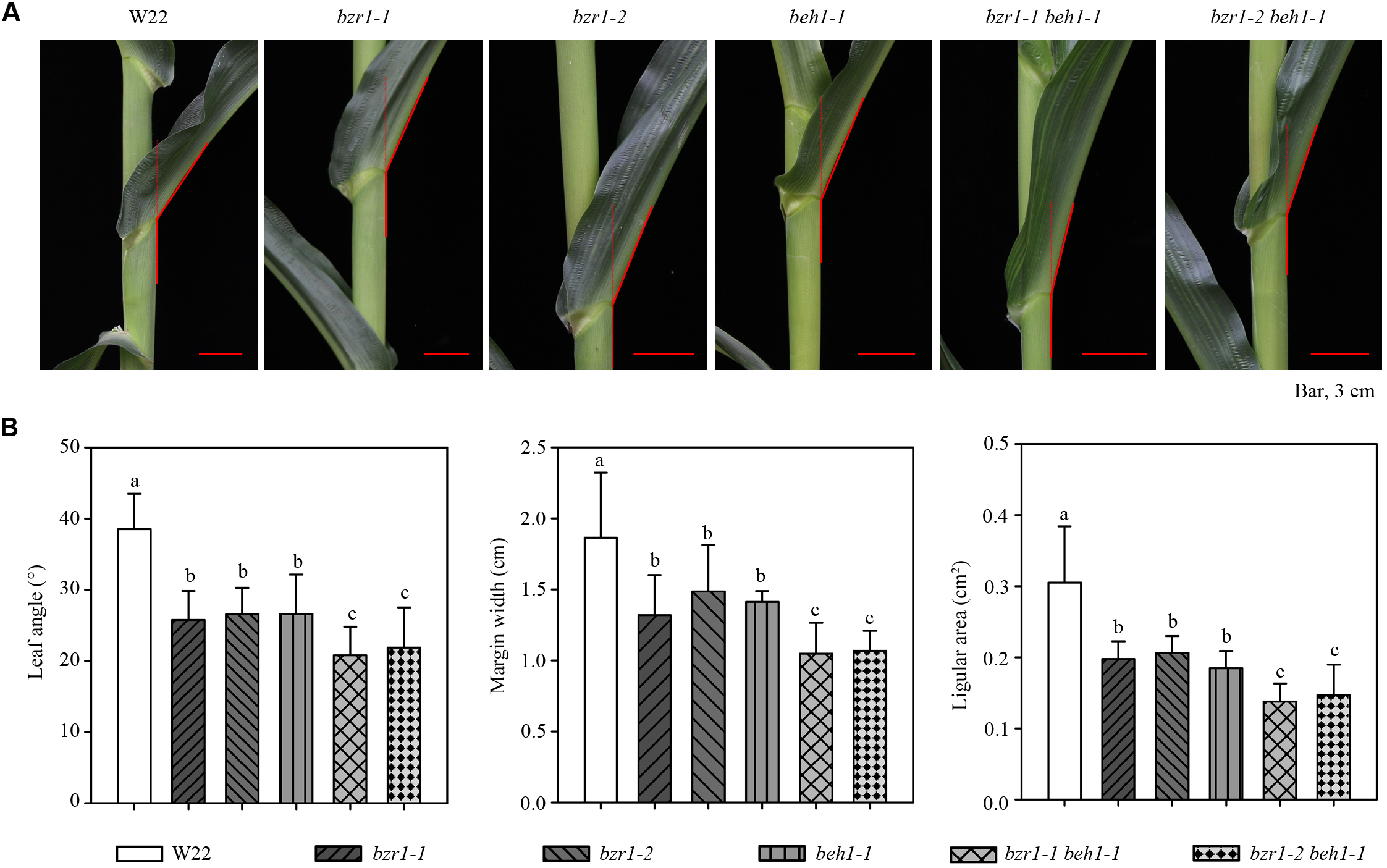
Leaf angle observation from single and double mutants of *bzr1* and *beh1*. A, Leaf angle morphology of WT inbred line W22, single mutants of *bzr1-1, bzr1-2* and *beh1-1,* double mutants of *bzr1-1 beh1-1* and *bzr1-2 beh1-1.* B, Quantitative measurements of leaf angle (n = 20), margin width (n = 10) and ligular area (n = 10) in W22, single and double mutants. Different letters above the columns indicate statistically significant differences between groups.

### ZmLG1 regulates the expression of Z*mBZR1*

Since ZmLG2 directly regulates the expression of *ZmBEH1*, we wondered whether ZmLG2 could also regulate the *ZmBZR1* as well. However, there was no ZmLG2 ChIP-seq binding site in the *ZmBZR1* promoter, and we could not detect the binding of ZmLG2 in Y1H and dual-luciferase assay. Motif analysis of *ZmBZR1* promoter identified an GTAC element, which is the binding motif of well-known transcription factor ZmLG1 controlling leaf angle (Tian et al., 2019; Moreno et al., 1997).

We then tested whether ZmLG1 could regulate the expression of *ZmBZR1*. Both Y1H and EMSA assay confirmed a direct binding of ZmLG1 in *ZmBZR1* promoter (Fig. 6, A and B). Next, dual-LUC transient assay performed in maize protoplasts confirmed that the ZmLG1 could activate the expression of *ZmBZR1,* as the ZmLG2 protein significantly induced the expression of LUC driven by the *ZmBZR1* promoter (Fig. 6C). Furthermore, we conduct the transient expression assay in tobacco (*Nicotiana tabacum*) leaves. The −640 to −168 sequence in front of TSS of *ZmBZR1* promoter was used to drive the expression of *LUC* gene. The strong LUC signals were detected only when the ZmLG1 protein was co-injected with the *ZmBZR1* promoter further validate the activation of ZmLG1 on the expression of *ZmBZR1* (Fig. 6D).

**Figure 6.**
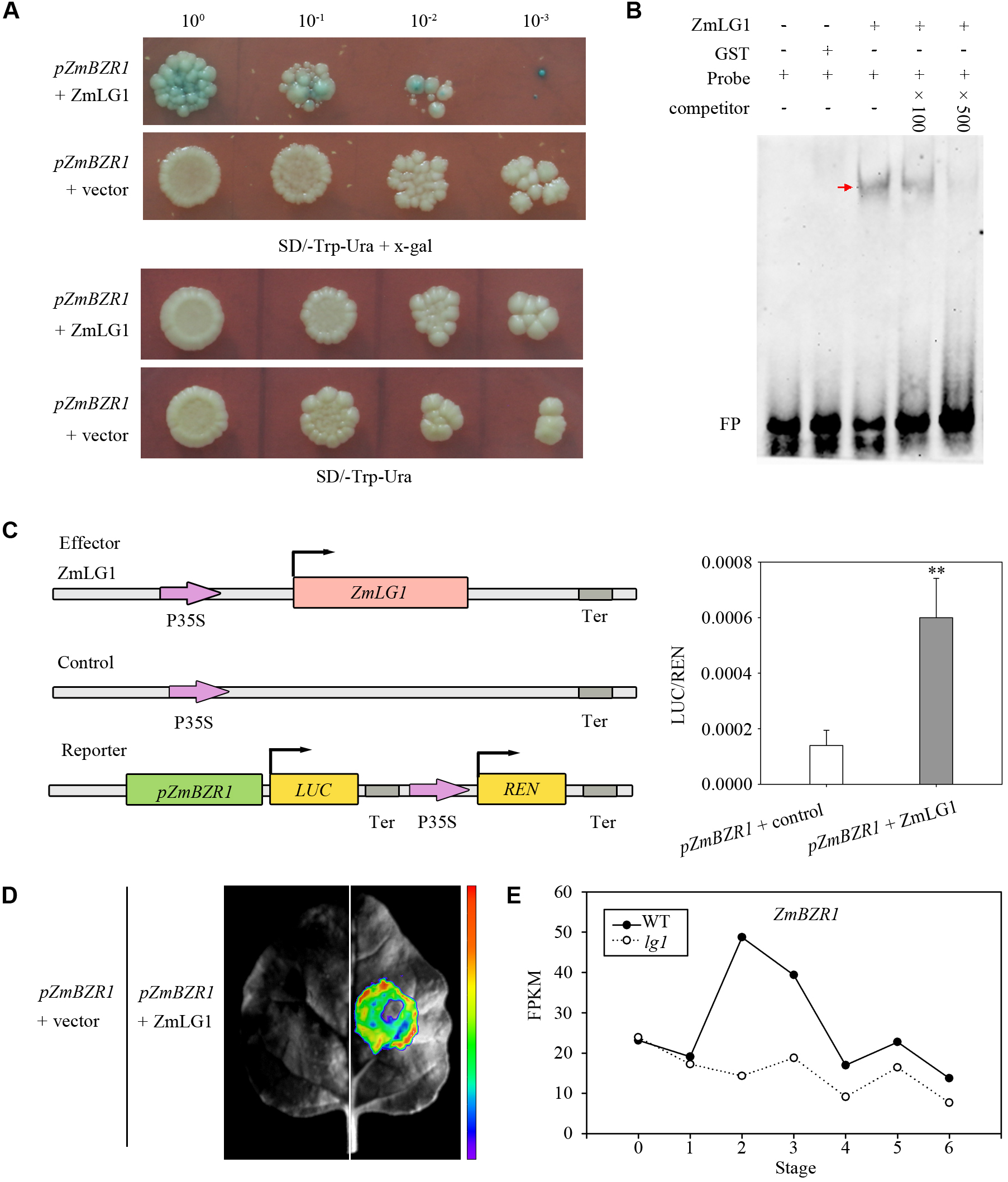
ZmLG1 directly activate the expression of *ZmBZR1*. A, ZmLG1 directly binds to the *ZmBZR1* promoter in yeast one hybrid (Y1H) assay. B, ZmLG1 directly binds to the *ZmBZR1* promoter in EMSA. “−” indicates the absence of proteins. “+” indicate the present; FP indicates free probe. C, ZmLG1 actives the promoter of *ZmBZR1* in Dual-luciferase assay. The coding sequence of LG1 driven by the 35S promoter was used as an effector, and the empty vector was used as an effector control in transient luciferase assay performed in maize protoplasts. The vector with the *Renilla reniformis* (REN) gene driven by a 35S promoter and firefly luciferase (LUC) gene driven by promoter sequence from *ZmBZR1* were used as the reporter. The LUC/REN ratio represents the relative activity of the promoters. Error bars represent SD (n = 5) **represents P < 0.01 determined by Student’s t test. D, Transient luciferase intensity assay performed in tobacco leaves showing that ZmLG1 actives the promoter of *ZmBZR1*. E, RNA-Seq data shows the expression of *ZmBZR1* in different maturation stage of ligular region (S0 to S4) in WT and *lg1* mutant. Details showed in Supplemental Fig. S2.

We also performed RNA-seq for the *lg1* mutant, and found that the expression of *ZmBZR1* was significantly reduced in the ligular region in *lg1* mutant than those in the WT (unpublished data, Supplemental Fig. S2B), especially in the middle stage of ligule expansion (stage 3 to 4), further indicate the positive regulation of ZmLG1 on the expression of *ZmBZR1* (Fig. 6E).

In addition, we generated the *lg1 bzr1-1* and *lg1 bzr1-2* double mutants to test the relationship between ZmLG1 and ZmBZR1, genetically. At the blade-sheath boundary, *lg1 bzr1-1* and *lg1 bzr1-2* have no ligule and auricle, which exhibit erect leaf architecture consistence with *lg1* (Supplemental Fig. S8), further indicates the epistatic effect of ZmLG1 on ligule and leaf angle formation.

### Yeast one-hybrid identified *ZmSCL28* as a downstream target of ZmBZR1 and ZmBEH1

To study how these transcription factors regulate leaf angle, we used yeast one-hybrid to find their potential downstream targets. Y1H assay showed that *ZmSCL28* promoter can be targeted by both ZmBZR1 and ZmBEH1 (Fig. 7A). ZmSCL28 encodes a GRAS domain transcription factor that has high homolog with rice *DLT* (Supplemental Fig. S9). In rice, OsBZR1 binds to CGTGCG elements (named BRRE motif) in the promoter of *DLT* gene to suppress its expression, and the similar regulation has also been reported in *Arabidopsis* (Li and Jin, 2007; Tong et al., 2009).

**Figure7.**
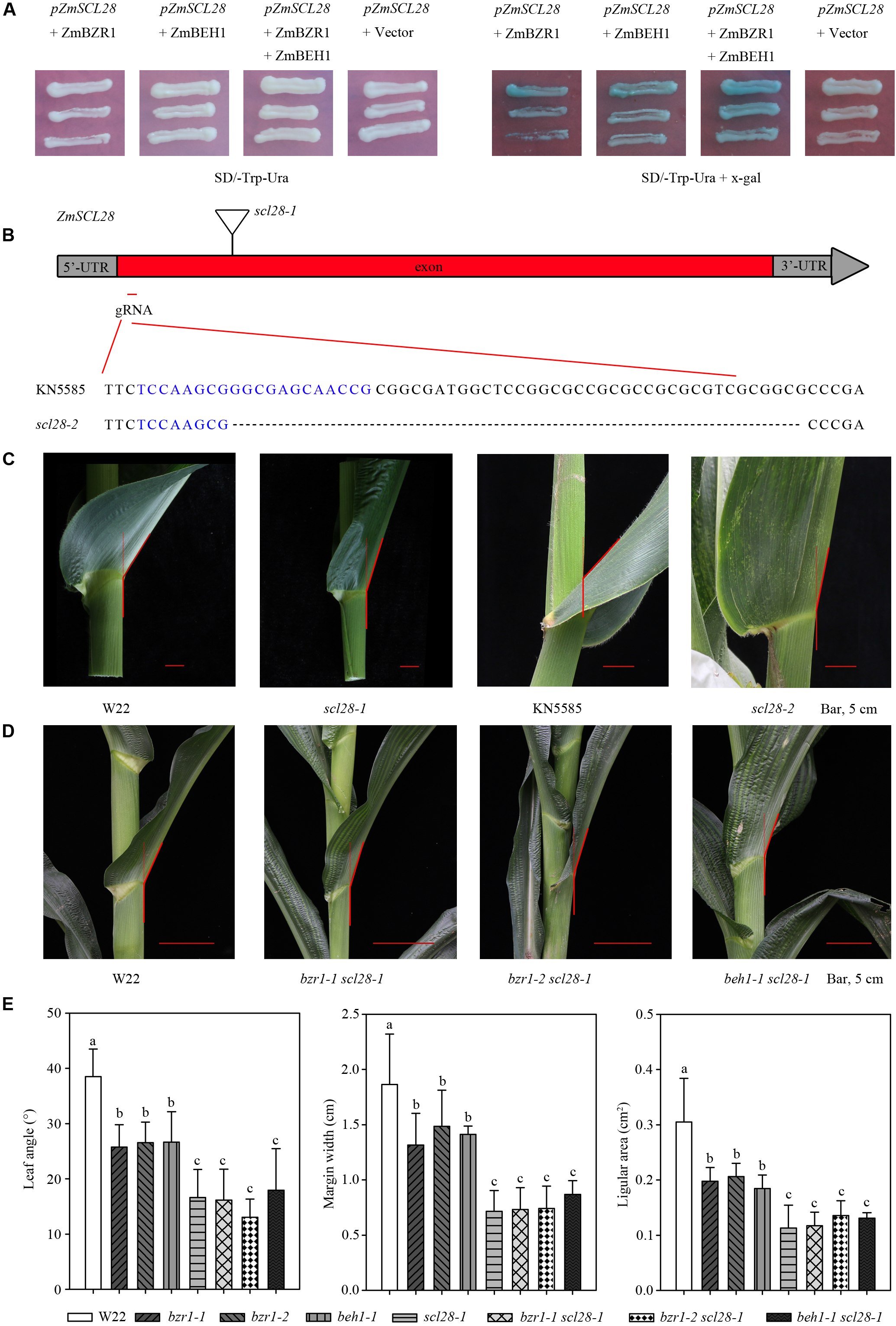
*ZmSCL28* works downstream of ZmBZR1 and ZmBEH1. A, Y1H assays showed that ZmBZR1 and ZmBEH1 bound to the promoter of *ZmSCL28*. B, Gene model presents the positions of two mutant alleles in *ZmSCL28*. The triangle shows the position of a Mu insertion (UFMu-09491) from line *scl28-1* at W22 genetic background, and red line shows the position of a single-guide RNA (sgRNA) designed to mutate the *ZmSCL28* using the CRISPR/Cas9 technology. The sequence of sgRNA is highlighted in blue, and the symbols “…” indicate the deletion caused by CRISPR/Cas9-induced mutations in line *scl28-2* at KN5585 genetic background. C, Leaf angle morphology in the WT inbred line W22 and mutant *scl28*-1, WT inbred line KN5585 and mutant *scl28-2* at V10 stage. Scale bars = 5 cm. D, Leaf angle morphology in the WT W22 and double mutants of *bzr1-1 scl28*-1, *bzr1-2 scl28*-1 and *beh1-1 scl28*-1 at V10 stage. Scale bars = 5 cm. E, Quantitative measurements of leaf angle (n = 20), margin width (n = 10) and ligular area (n = 10) in single and double mutants. Different letters above the columns indicate statistically significant differences between groups.

Next, we obtained a *ZmSCL28* uniform Mu insertion line (*scl28-1*) and confirmed the insertion in the first exon by PCR (Fig. 7B; Supplemental Fig. S3). The plant architecture of *scl28-1* is similar to that of *bzr1-1*, *bzr1-2* and *beh1-1*, conferring erect leaves, short margin width, small auricle size and decreased plant height, but the leaf angle is even smaller (Fig. 7, C, D and E; Supplemental Fig. S10, A and C).

To further confirm this, we generated a second loss-of-function mutant of *SCL28* by CRISPR/Cas9 (Fig. 7B). This *scl28-2* line has a 49 bp deletion located 50 bp downstream of the start codon. The phenotype of *scl28-2* is similar to *scl28-1*, *e.g.,* small leaf angle, upright leaf and short plant height (Fig. 7, C, D and E; Supplemental Fig. S10, A and C), confirming that *ZmSCL28* also plays important roles in controlling plant architecture.

To test the relationship among *ZmSCL28*, *ZmBZR1* and *ZmBEH1* genetically, the double mutants of *bzr1-1 scl28-1*, *bzr1-2 scl28-1* and *beh1-1 scl28-1* were generated. The double mutants had essentially the same phenotype as the *scl28-1* single mutant regarding plant height, leaf angle, auricle size, leaf length and width, with an exception being that the leaves of *bzr1-1 scl28-1* were slightly wider than that of *scl28-1* (Fig. 7, D and E; Supplemental Fig. S10, B and C). All these results support the conclusion that *ZmSCL28* is the direct target of ZmBZR1 and ZmBEH1 to regulate the leaf angle in maize.

## DISCUSSION

Increasing planting density is one of the key strategies to increase crop yield, and the selection of small leaf angle that determine the upright leaf architecture contribute significantly to the dense planting in maize (Tian et al., 2011; Pan et al., 2017; Wei et al., 2018; Tian et al., 2019; Zhao et al., 2019). ZmLG1 and ZmLG2 are two key loci that control the ligule development and then influence leaf angle formation, however, mutations at these two loci cause the abolish of normal ligule, which makes it easier for water to permeate into the leaf sheath, causing bacterial and fungal infections. In addition, the extremely erect leaf angle from the introgression lines of these alleles also limits their use in the commercial crop production. Therefore, we believe it is important to explore the downstream targets of ZmLG1 and ZmLG2 to achieve both small leaf angle and a normal ligule development.

To do so, we first examined the target of ZmLG2 and found *ZmBEH1*, a BZR1/BES1 homolog, and validate its role in controlling plant architecture. Our RNA-seq studies showed that ZmLG2 could regulate leaf angle through *ZmBEH1* in the stage of ligule/auricle expansion since the expression of *ZmBEH1* peaked at this stage (Fig.1D). The loss of function of *ZmBEH1* exhibit semi-dwarf phenotype, and the leaf angle was significantly decreased (Fig. 2, C and D). This decreasing may be caused by the reduced number of sclerenchyma cell layers at the adaxial site of the ligular region (Fig. 2F). However, how *ZmBEH1* regulates the sclerenchyma cell development on adaxial /abaxial sites needs further exploration.

Interestingly, Y2H and split LUC assay showed that ZmBEH1 could physically interact with ZmBZR1. The function of BZR1 has been well studied in *Arabidopsis* and rice, which plays central roles in brassinosteroid signaling pathways to affect plant development (i.e., erect leaves, phloem, and xylem differentiation), abiotic stress and so on (Wang et al., 2012; Sun et al., 2015; Saito et al., 2018; Min et al., 2019). The *ZmBZR1* mutant phenotypes were similar to that of *ZmBEH1*, e.g., semi-dwarf and decreased leaf angle (Fig. 4), however, the function of these two BES1/BZR1 family numbers were not redundantly at least in regulating the leaf angle formation, since the bzr*1-1 beh1-1* and *bzr1-2 beh1-1* double mutants displays smaller leaf angle than any of the single mutant (Fig. 5). We also found that ZmLG1, but not ZmLG2, could directly bind to the *ZmBZR1* promoter and active its expression (Fig. 6). The distinct regulation of ZmLG1 and ZmLG2 on *ZmBZR1* and *ZmBEH1* indicates a complex regulation specificity of ZmLG1 and ZmLG2 on the BR signaling factors, which further controls the maize leaf angle formation.

Next, we identified *ZmSCL28* as a direct downstream target of ZmBZR1 and ZmBEH1 through Y1H assay. *ZmSCL28* is the homolog gene of *DLT* in rice, the mutation of *ZmSCL28* shared similar phenotypes with *dlt*, e.g. dwarf and erect leaves, indicating the conserved function of these TFs in the different species. In rice, *DLT* was bind by OsBZR1 in the promoter through the BR-response element (Tong et al., 2009; Tong and Chu, 2009; Tong et al., 2012; Hirano et al., 2017; Qiao et al., 2017) and the loss-of-function mutant *dlt* was similar to BR-insensitive mutants (Tong et al., 2009). In maize, not only ZmBZR1, but also ZmBEH1 could bind to the BR-response element in the *ZmSCL28*. We also proved that *ZmSCL28* is the downstream target of ZmBZR1 and ZmBEH1 genetically, since the double mutants of *bzr1-1 scl28-1*, *bzr1-2 scl28-1* and *beh1-1 scl28-1* have almost identical phenotypes with *scl28-1* single mutant, for example, the leaf angle was 16.6° in *scl28-1* mutant and 16.1° in *bzr1-1 scl28-1* double mutant, however, the angle size was much larger in that of single mutants of *bzr1-1* (25.7°), *bzr1-2* (26.5°) and *beh-1* (26.6°) (Fig. 7).

Taken together, we propose a ZmLG2-BEH1-SCL28 regulatory cascade that effects leaf angle formation in maize (Fig. 8). In this model, we showed that *ZmBEH1* is directly activated by the ZmLG2, and ZmBEH1 can interact with ZmBZR1. *ZmBZR1* is directly activated by the ZmLG1, another well-known TF plays important roles in determine leaf architecture. Furthermore, we found that both ZmBEH1 and ZmBZR1 bind to the *ZmSCL28* promoter to regulate leaf angle. As the downstream targets of ZmLG1 or ZmLG2, the loss of function mutants of *ZmBEH1*, *ZmBZR1*, *ZmSCL28* have normal development of leaf ligule as expected, however, the size of leaf angle was dramatically reduced. Therefore, we propose that these three novel target loci, *ZmBZR1*, *ZmBEH1* and *ZmSCL28* are ideal targets to manipulate leaf angle in order to generate upright and semi-dwarf plant architecture, the two important traits pursued by breeders to increase the planting density and yields.

**Figure 8.**
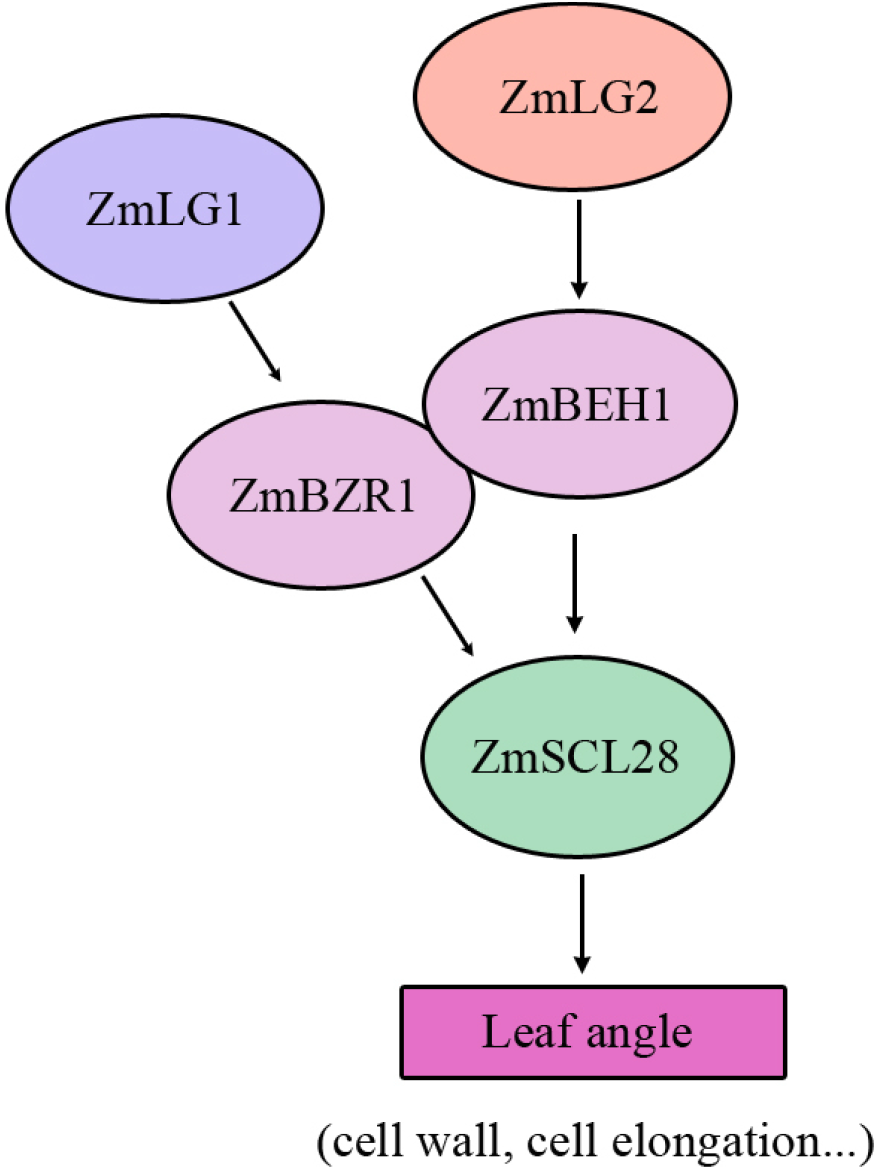
A proposed ZmLG2 (LG1)-ZmBEH1 (BZR1)-ZmSCL28 module regulates leaf angle in maize. The ZmLG2 binds to the protomer of *ZmBEH1* to activates its expression. The ZmBEH1 interact with ZmBZR1, which is a direct target of ZmLG1, to regulate the expression of a downstream target gene, *ZmSCL28*. These proposed regulatory cascades may control the leaf angle size in maize.

## Materials and Methods

### Plant Materials and Growth Conditions

Uniform Mu mutants with the stock number UFMu-13537, UFMu-03258, UFMu-13557 were requested from the maize genetic cooperation stock center. After two generations of backcrossing to maize inbred lines W22, the BC_2_F_1_ population that segregating the WT and mutant phenotypes were obtained. The stable single mutants from each gene were crossed to each other to obtain the double mutant.

To observe the phenotypes in seedling stage, seeds were planted into soil and were grown in a growth room with the setting of 28℃, 12-h-light/22℃, 12-h-dark cycle and 60% relative humidity.

### Yeast one-hybrid

To test the binding of ZmLG1 and ZmLG2 to the promoter of *ZmBZR1* or *ZmBEH1* in the yeast one-hybrid assay (Y1H), the full-length CDS of *ZmLG1*, *ZmLG2* were amplified and cloned into the pJG4-5 vector respectively to generate the destination plasmids. The fusion plasmids carrying the target genes were co-transformed with the *Lac Z* reporter vector (LacZi-2μ) driven by the promoter of *ZmBZR1* or *ZmBEH1* into the yeast strain EGY48. The empty vector pJG4-5 and the *Lac Z* reporter driven by the promoter of *ZmBZR1* or *ZmBEH1* were co-transformed as negative control.

Similarly, to test the binding of ZmBZR1 and ZmBEH1 to the *ZmSCL28* promoter, the full-length CDS of *ZmBZR1*, *ZmBEH1* were amplified and cloned into the pJG4-5 vector and the fusion plasmids were co-transformed with the *LacZ* reporter vector (LacZi-2μ) driven by the *ZmSCL28* promoter. The empty vector pJG4-5 and the *LacZ* reporter driven by the *ZmSCL28* promoter served as negative control.

The transformed yeast stains were plated on SD/-Trp-Ura medium and grown at 30°C for 3 days. After that, the transformants were tested on SD screening medium containing X-gal (5-bromo-4-chloro-3-indolyl-b-D-galactopyranoside) at 30°C for 4 days. The sequences of primers used in Y1H were listed in the Supplemental Table 1.

### Transient expression assay in maize protoplast

For the dual-luciferase transient expression assay, *ZmBZR1* and *ZmBEH1* promoter segments (808 bp and 1290 bp, respectively) were amplified from inbred line B73 and recombined into pGreenII 0800-LUC vector to generate the pZmBZR1/BEH1::LUC and pZmSCL28::LUC plasmids as reporter. The renillia luciferase (REN) gene driven by 35S promoter in the pGreenII 0800-LUC vector was used as the internal control. The full-length CDS of ZmLG1, ZmLG2 were amplified and recombined into pHW-3xAvi vector via Gateway LR Clonase II Enzyme mix driven by the 35S promoter, forming the effectors. The empty pHW-3 × Avi vector was used as control. The transient dual-luciferase assays were performed in the maize protoplasts collected from the leaves of 12-day-old etiolated seedling of inbred line B73. The luciferase signal was detected using dual-luciferase assay reagents following the manufacturer’s instructions. Relative LUC activity was calculated by normalizing LUC activity to REN activity. The sequences of primers used in the transient expression assay are listed in Supplementary Table S1.

### Transient expression assay in *Nicotiana benthamiana* leaves

The luciferase intensity assay was performed to examine the interactions between protein and promoter sequence. The full-length CDS of ZmLG1/LG2 and ZmBZR1/BEH1 was amplified and recombined into pCAMBIA1300-35s-GFP forming the effectors. *ZmSCL28* promoter (1048 bp) also recombined into pGreenII 0800-LUC vector with minimal CaMV 35S promoter. Then these constructs were transformed into the *Agrobacterium* strain GV3101. Next, Agrobacteria was infiltrated alone or together into the leaves of tobacco plants. About 3 d after infiltration, the transformed tobacco leaves were observed by Nightshade LB 985.All images were generated from LB985-Berthold.

### Electrophoretic Mobility Shift (EMSA) Assay

EMSA assay was performed as described previously (Song et al., 2020). The protein fused with GST has been purified firstly. The full-length BZR1/BEH1 cDNA was amplified with gene-specific primers and cloned into expression vector Pcold-GST. The construct was transformed into *Escherichia coli* BL21 cells, grown at 37°C. At OD_600_ = 0.4-0.5, the culture solution was placed at 16°C and let stand for 30 minutes. Isopropylthio-β-galactoside was added at the final concentration of 1 mM, and the culture was incubated at 16°C for 24 hours. The Pcold-GST-BZR1/BEH1 expressed GST-BZR1/BEH1 fusion protein was purified with BeaverBeads GSH (Beaver, Suzhou, China) and used for EMSA. The probes with biotin were synthesized according to manufacturer’s recommendation (Supplementary Table S1). The affinity bands labeled with biotin probes were detected by EMSA kit (Beyotime Biotechnology) and approximately 10 nm of probe was used for each binding assay. For competition assay, unlabeled probe was added to the reactions to detect the binding specificity.

### Yeast two-hybrid assays

The Yeast two-hybrid assay was performed using the Matchmaker™ Gold Yeast Two-Hybrid System (Clontech). The full-length coding region of *ZmBZR1* were ligated into pGBKT7 plasmid as bait and the full-length coding regions of *ZmBEH1* was fused into pGADT7 vector as prey. The bait and prey constructs were co-transformed into Y2H Gold yeast strain and grow at 30°C for 3 days. Next, the transformants were tested on SD screening medium containing X-α-Gal (5-Bromo-4-chloro-3-indoxyl-α-D-galactopyranoside) at 30°C for 4 d. Empty vectors were co-transformed as negative controls. The sequences of primers used in Y2H were listed in the Supplemental Table 1.

### The split luciferase complementation assay

The split firefly luciferase complementation assay was performed to examine the interactions between ZmBEH1 and ZmBZR1 using the constructs nLUC and cLUC. The full-length CDS of *ZmBZR1* without the stop codon and *ZmBEH1* with the stop codon were amplified and cloned into nLUC and cLUC, respectively, forming the nLUC-ZmBEH1 and ZmBZR1-cLUC constructs. *Agrobacterium* strain GV3101 cells carrying all constructs were transiently infiltrated into *Nicotiana benthamiana* leaves. After culturing for three days, luciferin (1 mM) was sprayed to activate luciferase, and the fluorescence signals were observed by the Chemiluminescent Imaging System (Tanon-5200). The primers used in the split firefly luciferase complementation assays are listed in Supplemental Table 1.

### Phylogenetic analysis of ZmBZR1/BEH1 and ZmSCL28

The full-length amino acid sequences of ZmBZR1/BEH1 and ZmSCL28 were used to BLAST search in protein databases of GRAMENE (http://www.gramene.org/), TAIR (http://www.arabidopsis.org) and RICEDATE (http://www.ricedata.cn/gene/) to identify the homologous genes in maize, *Arabidopsis* and rice. The amino acid sequences were aligned using the clustalw-2.0.10 software. A phylogenetic tree was constructed based on this alignment result using the neighbour-joining method in MEGA version 6 with the following parameters: Poisson correction, pairwise deletion, uniform rates and bootstrap (1,000 replicates).

### Statistical Analysis

To determine statistical significance, we employed Student’s test. (*P< 0.05, **P<0.01)

### RNA extraction and qPCR analysis

Total RNA was extracted using TRIzol reagent (Ambion) from maize leaves. The RNA samples were treated with DNase I (Thermo) and the concentration was measured by DS-11. Subsequently, cDNA was prepared using M-MLV Reverse Transcriptase and qRT-PCR were conducted using the SYBR Premix Ex Taq kit (Trans) on a Step One System (Applied Biosystems). The quantification method (2^−ΔCt^) was used and the gene expression level was estimated using three independent biological replicates. The maize *Ubi2* (UniProtKB/TrEMBL, Q42415) gene was used as an internal control to normalize the data. The process of qRT-PCR consisted of an initial denaturation step at 95 °C for 10 min, followed by 40 cycles at 95 °C for 15 s and 60 °C for 30 s. The primers for qRT-PCR are shown in Supplemental Table S1.

### Transformation

The CRISPR/Cas9 constructs for *ZmBEH1, ZmBZR1* and *ZmSCL28* were generated using a previously reported vector pBUE411 (Xing et al., 2014). The specific target sites and PAMs (the last three nucleotides) of *ZmBEH1, ZmBZR1* and *ZmSCL28* were designed and cloned into pBUE411 vector. These constructs were introduced into the *Agrobacterium tumefaciens* strain EHA105 and transformed into the maize inbred line KN5585 in WIMI Biotechnology Co., Ltd. The T_1_ plant was self-crossed to generate T_2_ plants for further research. The mutated sequences of T_2_ plants were confirmed by sequencing. The primers were shown in Supplementary Table S1.

## ACCESSION NUMBERS

Sequence data from this article can be found in Maize GDB Database under the following accession numbers: *ZmLG1* (*Zm00001d002005*), *ZmLG2* (*Zm00001d042777*), *ZmBZR1* (*Zm00001d021927*), *ZmBEH1* (*Zm00001d046305*), *ZmSCL28* (*Zm00001d045507*).

## ACKNOWLEDGMENTS

We thank the maize genetics cooperation stock center for kindly providing uniform Mu mutants with the stock number UFMu-13537, UFMu-03258, UFMu-13557. We thank Prof. Sarah Hake (University of California, Berkeley) for offering seeds of *lg1* and *lg2* mutants. We would also like to thank Prof. Feng Tian (China Agricultural University) for sharing the construct to induce the ZmLG1 recombinant proteins and Prof. Qijun Chen (China Agricultural University) for sharing the plasmid pBUE411.

---

**Supplemental Table S1.** The list of primers used in this paper.

**Supplemental Fig. S1.** Phylogenetic analysis of BZR1/BES1 family proteins in Maize, Rice and Arabidopsis.

**Supplemental Fig. S2.** Tissues of the ligular region collected from five different maturation stages of *lg2* (A) or seven of *lg1* (B).

**Supplemental Fig. S3.** PCR identification of *beh1-1, bzr1-1, bzr1-2* and *scl28-1* mutants.

**Supplemental Fig. S4.** Phenotypes of *beh1-1* mutants.

**Supplemental Fig. S5.** Agronomic characters of *bzr1-1* mutant.

**Supplemental Fig. S6.** Characterization of the CRISPR/Cas9-mediated knock-out mutants of *ZmBZR1*.

**Supplemental Fig. S7.** Phenotypes of the single and double mutants of *bzr1-1, bzr1-2* and *beh1-1*.

**Supplemental Fig. S8.** Phenotypes of single mutants and *lg1 bzr1-1, lg1 bzr1-2* double mutant plants.

**Supplemental Fig. S9.** Phylogenetic analysis of SCL family.

**Supplemental Fig. S10.** Phenotypes of *scl28-1/scl28-2* and *bzr1-1 scl28-1*, *bzr1-2 scl28-1*, *beh1-1 scl28-1* double mutant plants at 54-day-old stage.

## Literature Cited

Bai MY, Zhang LY, Gampala SS, Zhu SW, Song WY, Chong K, Wang ZY (2007) Functions of OsBZR1 and 14-3-3 proteins in brassinosteroid signaling in rice. Proc Natl Acad Sci USA 104: 13839–13844

Bolduc N, O’Connor D, Moon J, Lewis M, Hake S (2012) How to pattern a leaf. Cold Spring Harb Symp Quant Biol 77: 47–51

Cao Y, Zeng H, Ku L, Ren Z, Han Y, Su H, Dou D, Liu H, Dong Y, Zhu F, Li T, Zhao Q, Chen Y (2020) ZmIBH1-1 regulates plant architecture in maize. J Exp Bot 71: 2943–2955

Chen LG, Gao Z, Zhao Z, Liu X, Li Y, Zhang Y, Liu X, Sun Y, Tang W (2019) BZR1 family transcription factors function redundantly and indispensably in BR signaling but exhibit BRI1-independent function in regulating anther development in *Arabidopsis*. Mol Plant 12: 1408–1415

Duvick DN (2005) Genetic progress in yield of United States maize (*Zea mays* L.). Maydica: 193–202

Muehlbauer GJ, Fowler JE, Girard L, Tyers R, Harper L, Freeling M (1999) Ectopic expression of the maize homeobox gene *liguleless3* alters cell fates in the leaf. Plant Physiol 119: 651–662

Harper L, Freeling M (1996) Interactions of *liguleless1* and *liguleless2* function during ligule induction in maize. Genetics 144: 1871–1882

Hirano K, Yoshida H, Aya K, Kawamura M, Hayashi M, Hobo T, Sato-Izawa K, Kitano H, Ueguchi-Tanaka M, Matsuoka M (2017) Small organ size 1 and small organ size 2/dwarf and low tillering form a complex to integrate auxin and brassinosteroid signaling in rice. Mol Plant 10: 590–604

Kir G, Ye H, Nelissen H, Neelakandan AK, Kusnandar AS, Luo A, Inzé D, Sylvester AW, Yin Y, Becraft PW (2015) RNA interference knockdown of BRASSINOSTEROID INSENSITIVE1 in maize reveals novel functions for brassinosteroid signaling in controlling plant architecture. Plant Physiol 169: 826–839

Ku L, Wei X, Zhang S, Zhang J, Guo S, Chen Y (2011) Cloning and characterization of a putative *TAC1* ortholog associated with leaf angle in maize (*Zea mays* L.). PLoS ONE 6: e20621

Lambert RJ, Johnson RR (1978) Leaf angle, tassel morphology, and the performance of maize hybrids. Crop Sci 18: 499–502

Li H, Ye K, Shi Y, Cheng J, Zhang X, Yang S (2017) BZR1 Positively Regulates Freezing Tolerance via CBF-Dependent and CBF-Independent Pathways in *Arabidopsis*. Mol Plant 10: 545–559

Li JM, Jin H (2007) Regulation of brassinosteroid signaling. Trends Plant Sci 12: 37–41

Luo XM, Lin WH, Zhu S, Zhu JY, Sun Y, Fan XY, Cheng M, Hao Y, Oh E, Tian M, Liu L, Zhang M, Xie Q, Chong K, Wang ZY (2010) Integration of light- and brassinosteroid-signaling pathways by a GATA transcription factor in *Arabidopsis*. Dev Cell 19: 872–883

Makarevitch I, Thompson A, Muehlbauer GJ, Springer NM (2012) *Brd1* gene in maize encodes a brassinosteroid C-6 oxidase. PLoS ONE 7: e30798.

Mantilla-Perez MB, Salas Fernandez MG (2017) Differential manipulation of leaf angle throughout the canopy: current status and prospects. J Exp Bot 68: 5699–5717

Moreno MA, Harper LC, Krueger RW, Dellaporta SL, Freeling Michael (1997) *liguleless1* encodes a nuclear-localized protein required for induction of ligules and auricles during maize leaf organogenesis. Genes Dev: 616–628

Min HJ, Cui LH, Oh TR, Kim JH, Kim TW, Kim WT (2019) OsBZR1 turnover mediated by OsSK22-regulated U-box E3 ligase OsPUB24 in rice BR response. Plant J 99: 426–438

Moreno MA, Harper LC, Krueger RW, Dellaporta SL, Freeling M (1997) *liguleless1* encodes a nuclear-localized protein required for induction of ligules and auricles during maize leaf organogenesis. Genes Dev 11: 616–628

Pendleton JW, Smith GE, Winter SR, Johnston TJ (1968) Field investigations of the relationships of leaf angle in corn (*Zea mays* L.) to grain yield and apparent photosynthesis. Agronomy Journal 60: 422–424

Qiao S, Sun S, Wang L, et al. (2017) The RLA1/SMOS1 transcription factor functions with OsBZR1 to regulate brassinosteroid signaling and rice architecture. Plant Cell 29: 292–309

Ren Z, Wu L, Ku L, Wang H, Zeng H, Su H, Wei L, Dou D, Liu H, Cao Y, Zhang D, Han S, Chen Y (2020) *ZmILI1* regulates leaf angle by directly affecting *liguleless1* expression in maize. Plant Biotechnol J 18: 881–883

Saito M, Kondo Y, Fukuda H (2018) BES1 and BZR1 redundantly promote phloem and xylem differentiation. Plant Cell Physiol 59: 590–600

Song Z, Yan T, Liu J, Bian Y, Heng Y, Lin F, Jiang Y, Wang Deng X, Xu D (2020) BBX28/BBX29, HY5 and BBX30/31 form a feedback loop to fine-tune photomorphogenic development. Plant J 104: 377–390

Sun S, Chen D, Li X, Qiao S, Shi C, Li C, Shen H, Wang X (2015) Brassinosteroid signaling regulates leaf erectness in *Oryza sativa* via the control of a specific U-type cyclin and cell proliferation. Dev Cell 34: 220–228

Tian J, Wang C, Xia J, Wu L, Xu G, Wu W, Li D, Qin W, Han X, Chen Q, Jin W, Tian F (2019) Teosinte ligule allele narrows plant architecture and enhances high-density maize yields. Science 365: 658–664

Tong H, Chu C (2009) Roles of DLT in fine modulation on brassinosteroid response in rice. Plant Signal Behav 4: 438–439

Tong H, Jin Y, Liu W, Li F, Fang J, Yin Y, Qian Q, Zhu L, Chu C (2009) DWARF AND LOW-TILLERING, a new member of the GRAS family, plays positive roles in brassinosteroid signaling in rice. Plant J 58: 803–816

Tong H, Liu L, Jin Y, Du L, Yin Y, Qian Q, Zhu L, Chu C (2012) DWARF AND LOW-TILLERING acts as a direct downstream target of a GSK3/SHAGGY-like kinase to mediate brassinosteroid responses in rice. Plant Cell 24: 2562–2577

Tu X, Mejía-Guerra MK, Valdes Franco JA, Tzeng D, Chu PY, Shen W, Wei Y, Dai X, Li P, Buckler ES, Zhong S (2020) Reconstructing the maize leaf regulatory network using ChIP-seq data of 104 transcription factors. Nat Commun 11: 5089

Walsh J, Waters CA, Freeling M (1998) The maize gene *liguleless2* encodesa basic leucine zipper protein involvedin the establishment of the leaf blade-sheath boundary. Genes Dev 12: 208–218

Wang W, Bai MY, Wang ZY (2014) The brassinosteroid signaling network-a paradigm of signal integration. Curr Opin Plant Biol 21: 147–153

Wang ZY, Bai MY, Oh E, Zhu JY (2012) Brassinosteroid signaling network and regulation of photomorphogenesis. Annu Rev Genet 46: 701–724

Wei H, Zhao Y, Xie Y, Wang H (2018) Exploiting *SPL* genes to improve maize plant architecture tailored for high density planting. J Exp Bot 20: 4675–4688

Xing HL, Dong L, Wang ZP, Zhang HY, Han CY, Liu B, Wang XC, Chen QJ (2014) A CRISPR/Cas9 toolkit for multiplex genome editing in plants. BMC Plant Biol 14: 327

Zhang J, Ku LX, Han ZP, Guo SL, Liu HJ, Zhang ZZ, Cao LR, Cui XJ, Chen YH (2014) The *ZmCLA4* gene in the *qLA4-1* QTL controls leaf angle in maize (*Zea mays* L.). J Exp Bot 65: 5063–5076

Zhao Y, Wang H, Bo C, Dai W, Zhang X, Cai R, Gu L, Ma Q, Jiang H, Zhu J, Cheng B (2018) Genome-wide association study of maize plant architecture using F _1_ populations. Plant Mol Biol 99: 1–15

